# Invasive grass causes biotic homogenization in wetland birds

**DOI:** 10.1101/2021.07.03.451016

**Authors:** C.D. Robichaud, R.C. Rooney

**Affiliations:** B2-251, University of Waterloo, Department of Biology, 200 University Ave. West, N2L 3G1

**Keywords:** *Phragmites australis*, invasive species, bird community, wetland, coastal marsh, water-level fluctuations, biological homogenization, beta diversity

## Abstract

Plant invasions often lead to homogenization of the plant community, but the potential for plant invasions to cause homogenization of other trophic levels is under-studied in many systems. We tested whether the bird community in *Phragmites australis-*invaded marsh would exhibit spatial and temporal taxonomic homogenization compared to remnant cattail and meadow marsh. We compared the bird community using marsh invaded by *P. australis* and remnant, uninvaded marsh vegetation in a year with average water depths and a year with above-average water depths in the coastal marshes of a World Biosphere Reserve. Our results demonstrate strong evidence for spatial and temporal homogenization of the wetland bird community following *P. australis* invasion. The birds present in *P. australis*-invaded marsh were a nested subset of those present in remnant marsh, and total beta diversity decreased when water depths were above average. In contrast, total beta diversity was high in remnant marsh vegetation and stable between the two years. The distinctively structured vegetation zones in remnant (uninvaded) marsh yields structural complexity and habitat heterogeneity that supports greater taxonomic turnover in the bird community. Our study provides evidence that invasion by a plant has resulted in biological homogenization of the wetland bird community.

## Introduction

Globally, invasive species pose a major threat to biodiversity (IUCN 2020). One way in which invasive species alter ecosystems is biotic homogenization, where communities become more similar over time (McKinney and Lockwood 1999, Olden and Poff 2004). As a result, invasion creates communities that are taxonomically and functionally similar due to the shared presence of an abundant introduced species (McKinney and La Sorte 2007). This can lead to a loss of habitat heterogeneity, a property of ecosystems that is associated with high species diversity (e.g., Kadmon and Allouche 2007). Invasive plants can alter properties of ecosystems that are important for other taxa, such as vertical and horizontal structure (e.g., Gagnon Lupien et al. 2015), or nutrient cycling and productivity (e.g., Vilà et al. 2011). Thus, homogenization as a result of invasion can lead to the extirpation or extinction of native species (McKinney and Lockwood 1999).

Freshwater coastal marshes provide a model system for testing the homogenizing effect of invasive species. Coastal marshes are highly dynamic systems that are maintained by spatial and temporal heterogeneity in water depth. This is key, as vegetation communities are closely tied to hydrology such that there is zonation in accordance with variations in water depth (Keddy and Reznicek 1986, Keddy and Campbell 2019). As coastal marshes are defined by this heterogeneity, any homogenization promoted by biological invasion should affect communities adapted to marsh habitats, even over short timescales (e.g., Price et al. 2018, Muthukrishnan and Larkin 2020). This can be particularly detrimental to wetland species, as biological diversity in wetlands is already restricted to the small fraction of taxa that possess adaptations necessary to survive variable periods of inundation (e.g., Correll et al. 2016, Daniel et al. 2019). As such, marsh birds are an excellent candidate species to examine biotic homogenization in freshwater coastal marshes as their resources (e.g., prey, nesting sites) are closely tied to water depth fluctuations and many species have preferences for certain vegetation communities (Riffell et al. 2001, Desgranges et al. 2006). Any process that leads to the homogenization of wetlands is expected to reduce the diversity of bird communities (Steen et al. 2006).

The water level variations that are essential for coastal marshes also make them vulnerable to colonization by invasive plant species (Galatowitsch et al. 1999, Tulbure et al. 2007). Wetlands throughout North America are presently facing invasion by European Common Reed (*Phragmites australis* subsp. *australis* ([Cav.] Trin. ex Steud.). *Phragmites australis* is a widely distributed invasive wetland grass (Catling and Mitrow 2011) which creates monocultures (Packer et al. 2017) that displace resident vegetation (e.g. Wilcox et al. 2003, Robichaud and Rooney 2021), fill in open-water pools (Able and Hagan 2013), and alter wetland hydrology (Weinstein and Balletto 1999). For example, we observed that *P. australis*-invaded marsh exhibits less seasonal drawdown than uninvaded areas (Yuckin and Rooney 2019). While native marsh vegetation communities are restricted by environmental conditions, such as water depth and inundation period, *P. australis* can establish populations across a wide moisture gradient (Packer et al. 2017) and spans the entire water depth gradient occupied by resident wetland vegetation (Wilcox et al. 2003, Robichaud and Rooney 2021). The wide environmental tolerances of *P. australis* allow it to distrupt the organization of native vegetation, resulting in the loss of vegetation diversity and zonation in favour of near monocultures of dense, emergent grass. We expect that this homogenization of vegetation and hydrologic conditions would alter bird communities, but the documented responses to *P. australis* invasion are complex. In some marshes, there is little difference in the abundance or identity of birds using invaded and uninvaded habitat (e.g., Gagnon Lupien et al. 2015), while others have found a negative relationship between bird diversity and *P. australis* (e.g., Prosser et al. 2018) that may take decades to become evident (e.g., Robichaud and Rooney 2017). The spatial and temporal homogenization of both hydrologic and vegetation structure in *P. australis*-invaded marsh, likely also leads to homogenization of the marsh avifauna. Yet, no studies have directly tested the hypothesis that invasion by a wetland plant could lead to biotic homogenization of the bird community or that birds respond differently to year-to-year water depth fluctuations between *P. australis-*invaded and reference vegetation communities.

To fill this knowledge gap, we conducted a field experiment in the coastal marshes of Long Point, ON. We compared bird communities in *P. australis-*invaded marsh to those in remnants of uninvaded marsh vegetation in a year with average water depths and a year with above-average water depths. We tested two hypotheses: 1) that bird communities associated with *P. australis*-invaded marsh have lower beta diversity than in remnant marsh vegetation (spatial homogenization), and 2) that bird community diversity is less responsive to year-to-year water depth changes in *P. australis*-invaded marsh than in uninvaded, remnant marsh (temporal homogenization).

## Methods

### Study area and field methods

We sampled bird communities in Long Point, Ontario, CA ([42°32′51″N, 80°3′33″W](https://geohack.toolforge.org/geohack.php?pagename=Long_Point,_Ontario&params=42_32_51_N_80_3_33_W_region:CA-ON_type:city); Fig. 1), a sandspit peninsula that contains roughly 70% of the remaining intact coastal marsh on the north shore of Lake Erie (Ball et al. 2003). Long Point is a World Biosphere Reserve, Ramsar wetland of international importance, and Important Bird Area. In the late 1990s, invasive *P. australis* began to spread in these marshes and replaced resident meadow marsh and emergent marsh vegetation communities with monocultures of *P. australis* (Wilcox et al. 2003, Wilcox 2012). Meadow marsh and emergent marsh represent vegetation communities (hereafter ‘remnant marsh’) present on either end of the wetland moisture gradient in Long Point. Meadow marsh has saturated soils and periods of shallow standing water, leading to diverse vegetation communities consisting of grasses, sedges, and forbs. Emergent marsh, in contrast, has standing water all growing season and a vegetation community composed of robust emergent vegetation (e.g., *Typha* spp.). With its ability to tolerate a wide range of environmental conditions (Packer et al. 2017), *P. australis* replaces both emergent and meadow marsh communities, reducing their extent in Long Point (Wilcox et al. 2003, Jung et al. 2017).

**Figure 1.**
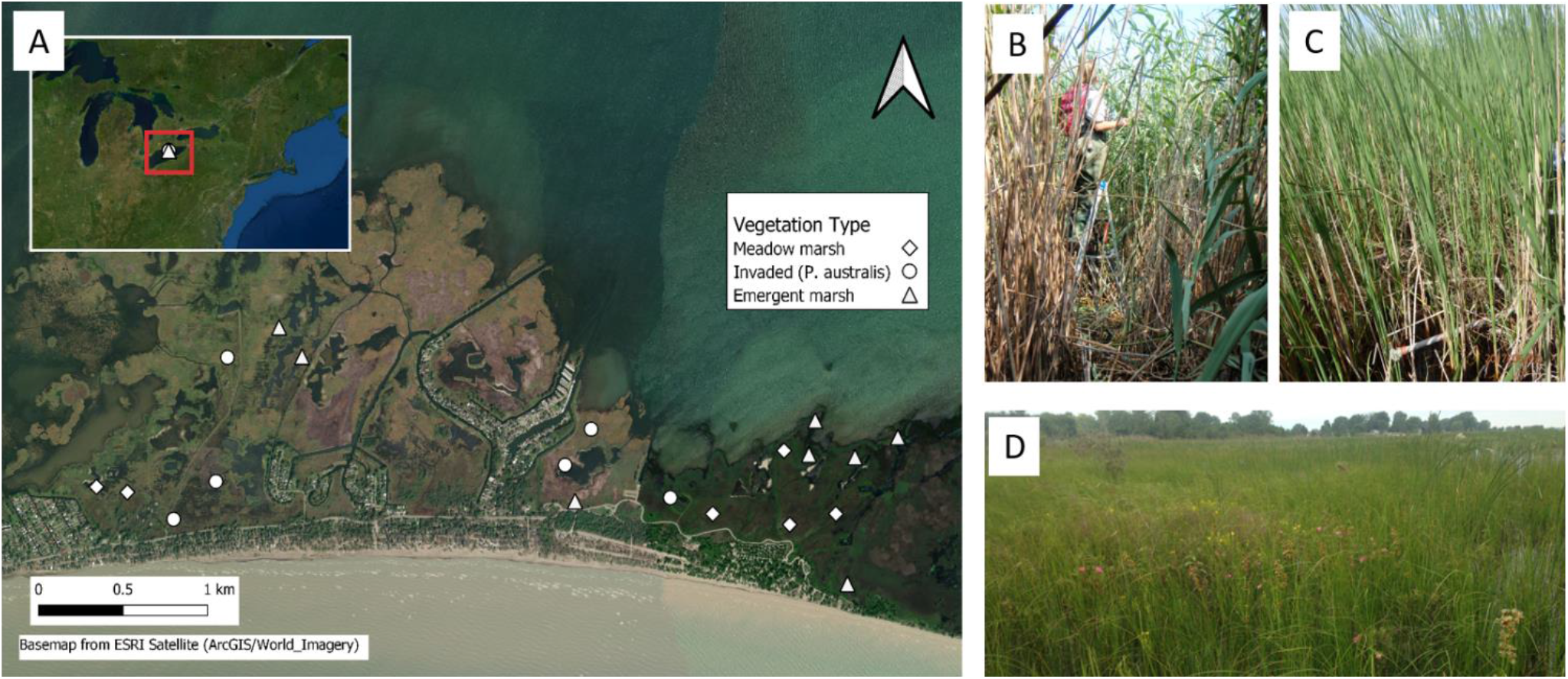
Bird survey point-counts that were established in 2014 and re-visited in 2015 in the western portion of Long Point peninsula, ON, Canada (A). Surveys were placed in areas invaded by *Phragmites australis* (n = 6) (B) and in remnant, uninvaded vegetation consisting of emergent marsh (n = 8) (C) and meadow marsh (n = 6) (D). Figure made with QGIS, base map from ESRI Satellite (ArcGIS/World_Imagery).

We surveyed birds in the western portion of Long Point by establishing twenty point-count locations in 2014 that we surveyed again in 2015; six in *P. australis-*invaded communities (“invaded”), six in remnant meadow marsh, and eight in remnant emergent (i.e. cattail) marsh (Fig. 1). We visited each point count four times during the breeding season, from early May to early August, separated by at least 10 days between visits. Point counts took place a half hour before sunrise to 09:30 with visits to each site randomized each day to reduce temporal bias. We modified the Marsh Monitoring Protocol (Bird Studies Canada 2009) to accommodate for the challenges of detection among marsh vegetation types. For all point counts, we used a 1.8 m ladder to standardize observer height and recorded all individual birds seen or heard within a 25 m radius around the center point of the survey or foraging 100 m above the point count (Meyer 2010). This reduced radius was used to standardize sampling effort among the different vegetation types and was based on previous work using broadcast recordings of eight common marsh birds to be the maximum radius across which all eight species were reliably detected in all three vegetation types (Meyer 2003).

Birds that flew straight through the survey point (“fly throughs”) were not recorded, so counts indicate birds that were actively using each site. Surveys were 15 minutes in length and consisted of 5 minutes of passive listening, 5 minutes of call broadcasting for secretive marsh birds (Virginia Rail (*Rallus limicola*), Sora (*Porzana carolina*), Least Bittern (*Ixobrychus exilis)*, a combination of Common Moorhen (*Gallinula chloropus*) and American Coot (*Fulica americana*), and Pied-billed Grebe (*Podilymbus podiceps*), and a final 5 minutes of passive listening (Bird Studies Canada 2009, Robichaud and Rooney 2017). The observer was the same for all point counts both years to reduce survey bias. In mid-August we measured water depth at each site. Water depths in Lake Erie exhibit a strong seasonal cycle, with peaks in June and lows in December, and daily water levels can fluctuate up to 50 cm due to seiche events (Farhadzadeh 2017). This daily and seasonal variation is superimposed on multi-decadal climate-driven cycles in lake depths (e.g., Holcombe et al. 2003, Wiles et al. 2009). Lake Erie water levels during the survey period were marginally higher than the historical average in 2014, and considerably higher in 2015, which was reflected at the site level (Appendix A).

### Statistical approach

#### Bird abundance and alpha diversity

Since *P. australis* replaces both meadow marsh and emergent marsh communities, we combined these communities into “remnant marsh” for analyses. Analyses with communities separated are available in the Appendices. Recognizing some birds will use the marsh at different times over the breeding season, we summed bird observations across all four visits each year to capture the variation in bird presence (Robichaud and Rooney 2017). To account for imperfect detections, we calculated the estimated asymptote for Hill numbers for species richness (q = 0), Shannon-Weiner diversity (q = 1), and Simpson’s diversity (q = 2) (Chao et al. 2014) for each vegetation type each year using the iNEXT package (Hsieh et al. 2020). To compare total abundance, species richness, Shannon-Weiner diversity, and Pielou’s evenness using a general linear model we also calculated these values for each site using the vegan package (Oksanen et al. 2019). We modeled total abundance, richness, Shannon-Weiner diversity, and Pielou’s evenness as a function of vegetation community type and year using a two-way ANOVA (Type III SS) with an interaction term. To improve the normality of the residuals, we log_10_ transformed total abundance and species richness data. Analyses were performed using the car package (Fox and Weisberg 2019) in R v 4.0.0 (R Core Team 2016).

### Community composition and beta diversity

To assess differences in beta diversity, or the compositional difference between sites (Anderson et al. 2011), between *P. australis*-invaded and remnant vegetation types, we measured the average dissimilarity of individual observations to their group centroid (i.e., themultivariate homogeneity of variances) using PERMDISP2 (Anderson et al. 2006). We repeated this assessment with remnant marsh broken into meadow marsh and emergent marsh vegetation categories to better parse out the variation in these two vegetation communities. To test if one group was more variable than another, we conducted a permutational test using the least-square residuals to generate an *F* distribution to assess the likelihood of no difference between groups (Anderson et al. 2006, Oksanen et al. 2019). Using a Bray-Curtis dissimilarity matrix, we ran the PERMDISP2 procedure and the permutation test, with 999 permutations, using the *betadisper* and *permutest*.*betadisper* functions from the vegan package (Oksanen et al. 2019). To visualize the bird community composition among the vegetation types we used a non-metric multidimensional scaling ordination. For the NMDS ordination, we used a Bray-Curtis dissimilarity matrix and first assessed the stress of solutions ranging from 1 to 10 axes to determine the best number of axes for the data. The optimal ordination solution was determined using an iterative process with a random starting configuration, and iterations stopped once two convergent solutions were achieved. The ordination was run with the *metaMDS* function from the vegan package (Oksanen et al. 2019).

Beta diversity can result from turnover, i.e., species replacement between sites, and/or nestedness, i.e., the exclusion of species from certain sites, such that the species at one site are a subset of species at another (Baselga 2010). To identify the processes that contribute to beta diversity in each vegetation community we computed beta diversity and its components using a presence/absence matrix to provide equal weight to all species. Using pair-wise Sørensen dissimilarity, we calculated beta diversity, turnover and nestedness for each vegetation type and year combination using the betapart package (Baselga and Orme 2012). We also generated null-models (Anderson et al. 2011) by pooling the bird species present in both years in either remnant or invaded vegetation communities, and then randomizing the resulting matrix while maintaining species occurrence frequency and sample species richness (Gotelli 2000) using the picante package (Kembel et al. 2010). We then split the randomized matrix by year and computed beta diversity, nestedness, and turnover for the vegetation and year combinations. We did this 1000 times, taking the average and 95% CI of beta diversity sum, turnover, and nestedness for comparison with our observed values.

To quantify the uniqueness and relative contribution to total beta diversity of each site, we calculated the local contribution to beta diversity (LCBD) using the methods specified in Legendre and De Cáceres (2013) and Kuczynski et al. (2018). Since LCBD values are standardized, we can compare values between years for each site (Kuczynski et al. 2018) by calculating the differences as LCBD_2015_ – LCBD_2014_, where a negative value indicates the site has become more homogenous between years and a positive value indicates differentiation. We used a Bray-Curtis dissimilarity matrix, so that differences in relative abundance were accounted for, and performed these analyses using the adespatial package (Dray et al. 2020).

As a complement to the beta diversity and LCBD analyses, we also quantified the changes in community composition between the two years at each site by calculating the temporal beta-diversity index (TBI) (Legendre 2019). TBI values range from 0 (exactly the same) to 1 (maximum dissimilarity) and can be decomposed into species gains (C) and species losses (B) by comparing the number of species at each time point. This analysis provides insight into why beta diversity, or total dissimilarity (D), changed between the two years (Kuczynski et al. 2018, Legendre 2019). We calculated TBI using Bray-Curtis dissimilarity in the adespatial package (Dray et al. 2020). We conducted all analyses using R v. 4.0.0. (R Core Team 2016).

## Results

### Bird abundance and alpha diversity

We observed a total of 23 bird species: 17 in 2014 and 20 in 2015. In 2014, we detected 14 species in remnant marsh and 12 in *P. australis-*invaded marsh. In 2015, we detected 20 species in remnant marsh and 8 in *P. australis-*invaded marsh. The asymptotic species richness estimated in *P. australis*-invaded marsh in 2014 (19.92) was lower than the richness estimated in remnant marsh (31.9) (Figure 2A; Table 1). In 2015, this difference increased as the species richness estimate dropped to 8.24 in *P. australis*-invaded marsh while remnant marsh had an estimate of 28.14 (Fig. 2B; Table 1). The richness estimates for emergent marsh remained similar in both years, but richness estimates increased in meadow marsh from 8.98 (8.09 – 18.91 (95% confidence limits)) to 21.13 (17.67 – 42.60 (95% confidence limits)) between 2014 and 2015 (Appendix B).

**Figure 2.**
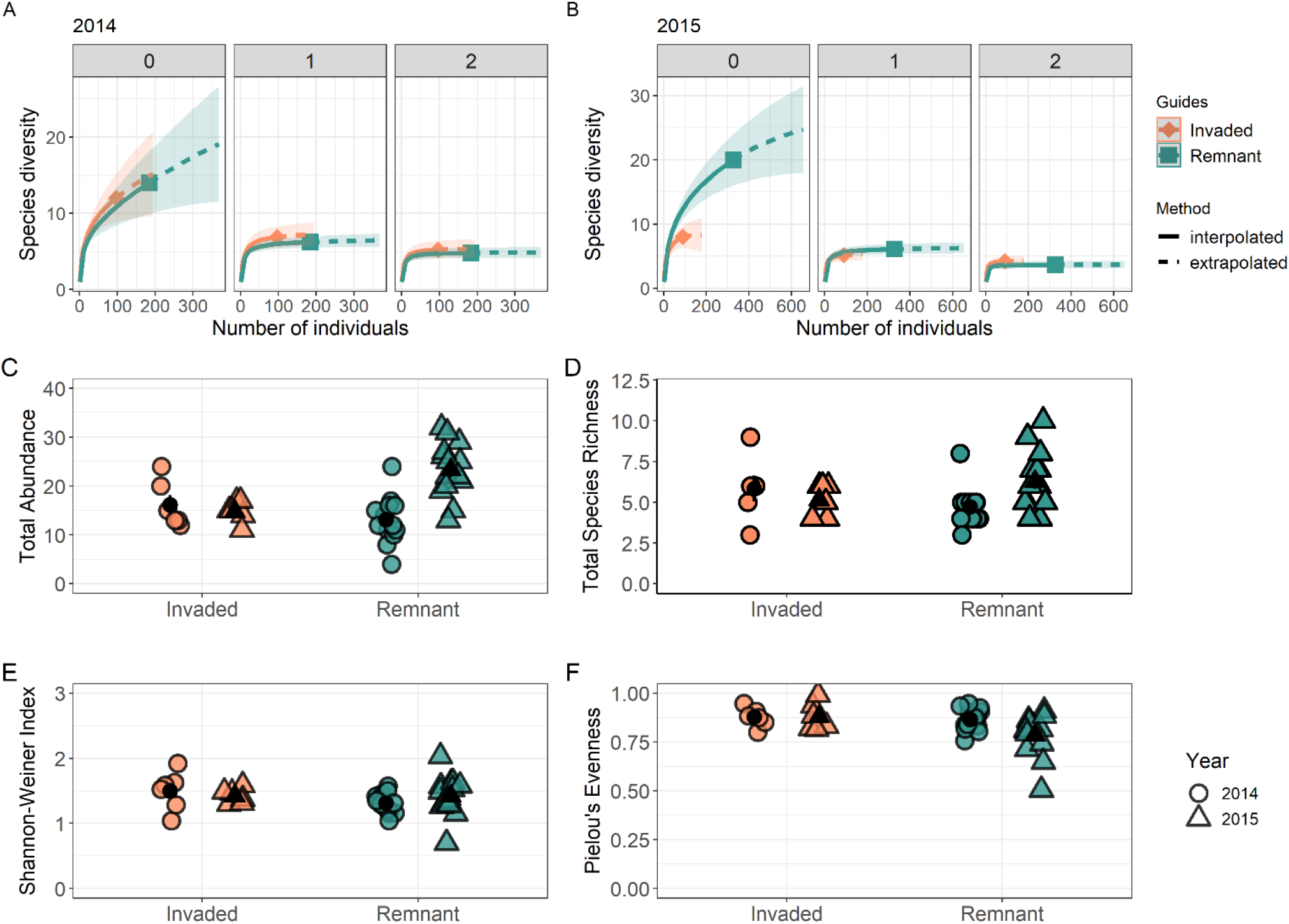
Sample-sized based rarefaction and extrapolation sampling curves for remnant and *P*. australis-invaded habitat in 2014 (A) and 2015 (B) by diversity order: 0 = species richness, 1 = Shannon-Weiner diversity, 2 = Simpson’s diversity. Points represent the reference samples and shaded areas represent 95% confidence intervals. The degree of change in total bird abundance (C), species richness (D), Shannon-Weiner index (E), and Pielou’s evenness (F) in remnant marsh and *P. australis* invaded marsh between 2014 and 2015. High Lake Erie water levels in 2015 resulted in deeper standing water in all vegetation communities. High abundances in 2015 are a result of remnant meadow marsh vegetation. Error bars represent standard error.

**Table 1.** Aysmptotic diversity estimates of species richness, Shannon-Weiner diversity, and Simpson’s diversity calculated using iNEXT. Values in parantheses are 95% upper and lower confidence intervals.

|  | Remnant vegetation |  | *P. australis*-invaded habitat |  |
| --- | --- | --- | --- | --- |
|  | 2014 | 2015 | 2014 | 2015 |
| Species richness | 31.90 (16.52-141.38) | 28.14 (21.57-62.30) | 19.92 (12.98-76.18) | 8.25 (8.01-12.69) |
| Shannon-Weiner | 1.90 (1.83-2.10) | 1.85 (1.81-2.01) | 2.03 (1.93-2.27) | 1.68 (1.64-1.87) |
| Simpson's diversity | 0.80 (0.79-0.82) | 0.73 (0.73-0.77) | 0.82 (0.81-0.86) | 0.77 (0.76-0.82) |

The degree of change in total bird abundance between 2014 and 2015 varied depending on which vegetation community was considered, as evidenced by significant vegetation type-year interactions for abundance (two-way ANOVA F_1,36_ = 8.78, p = 0.005; Fig. 2C). In remnant marsh, the average bird abundance was higher in the wetter 2015, whereas in *P. australis-*invaded sites there was little change between years. Metrics related to alpha diversity, including species richness, Shannon-Weiner index, and Pielou’s evenness (Table 2; Fig. 2D-F) did not significantly differ between vegetation types or year. Though not statistically significant, there was an increase in the species richness of birds in remnant marsh in 2015 (S_2014_ = 14, S_2015_ = 20, 43% increase), especially within remnant meadow marsh (S_2014_ = 8, S_2015_ = 17, 112% increase), compared to invaded marsh (S_2014_ = 11, S_2015_ = 8, 18% decrease) (Appendix C & D).

**Table 2.** Two-way ANOVAs (Type III SS) were used to assess if abundance, species richness, diversity or evenness of the bird community differed between vegetation types either year. Abundance and species richness were log transformed for analyses.

|  | Abundance |  |  | Species Richness |  | Shannon-Weiner diversity |  | Pielou's evenness |  |
| --- | --- | --- | --- | --- | --- | --- | --- | --- | --- |
|  | df | F | p | F | p | F | p | F | p |
| Vegetation | 1 | 2.451 | 0.126 | 2.094 | 0.157 | 2.546 | 0.119 | 0.089 | 0.767 |
| Year | 1 | 0.116 | 0.735 | 0.327 | 0.571 | 0.322 | 0.574 | 0.002 | 0.964 |
| Interaction | 1 | 9.483 | 0.004 | 3.922 | 0.055 | 1.265 | 0.268 | 2.277 | 0.140 |
| Residuals | 36 |  |  |  |  |  |  |  |  |

In both 2015 and 2014, Red-winged Blackbirds were most abundant in *P. australis* vegetation (66 total), followed by Swamp Sparrow (34 total), Marsh Wren (29 total), and Common Yellowthroat (28 total). In 2014, we detected two wetland obligates, Virginia Rail and Sora, and an American Woodcock in *P. australis*. In 2015, only eight species were detected in *P. australis*: Marsh Wren, Swamp Sparrow, Common Yellowthroat, Red-winged Blackbird, Song Sparrow, Yellow Warbler, Barn Swallow and Eastern Kingbird (Appendix C). Of these 8 species, 90% of the observed individuals were Marsh Wren, Swamp Sparrow, Red-winged Blackbird, or Common Yellowthroat.

Red-winged Blackbirds were also most abundant in both remnant meadow marsh vegetation (84 total) and in remnant emergent marsh vegetation, where their abundance increased from 39 total in 2014 to 94 total in 2015. Similar to *P. australis*, Swamp Sparrow (50 total), Common Yellowthroat (49 total), and Marsh Wren (51 total) were the most abundant species in cattail marsh both years. Wetland obligates American Bittern, Least Bittern, and Virginia Rail were detected in emergent marsh in both years. In remnant meadow marsh, Swamp Sparrow (22 total) and Common Yellowthroat (33 total) were very abundant both years, but Marsh Wren did not use meadow marsh habitat until 2015, when water levels were higher. Fewer wetland obligate species were detected in meadow marsh in 2014, though Virginia Rail and Sora were detected in 2015 (Appendix E).

### Community composition and beta diversity

*Phragmites australis-*invaded habitat had considerably lower beta diversity in 2015 compared to remnant vegetation both years (permutational pairwise comparisons: p ≤ 0.037, Appendix E). The average dissimilarity to the group centroid in invaded marsh was 0.505 and 0.371, in 2014 and 2015, respectively. Beta diversity was greater in remnant marsh, as the average dissimilarity to the group centroid was 0.528 and 0.542 in 2014 and 2015, respectively. When we broke remnant marsh into meadow marsh and emergent marsh, emergent marsh had a slight decrease in the average distance to the group centroid from 2014 (0.508) to 2015 (0.451) while meadow marsh had an increase distance to group centroid from 2014 (0.446) to 2015 (0.541) (Appendix F). The lower beta diversity in *P. australis-*invaded marsh is evident in the NMDS ordination (Fig. 3). The final ordination was a 3D configuration with a stress of 15.56 that found two convergent solutions after 20 attempts (non-metric fit, R^2^ = 0.976) of a maximum 1000.

**Figure 3.**
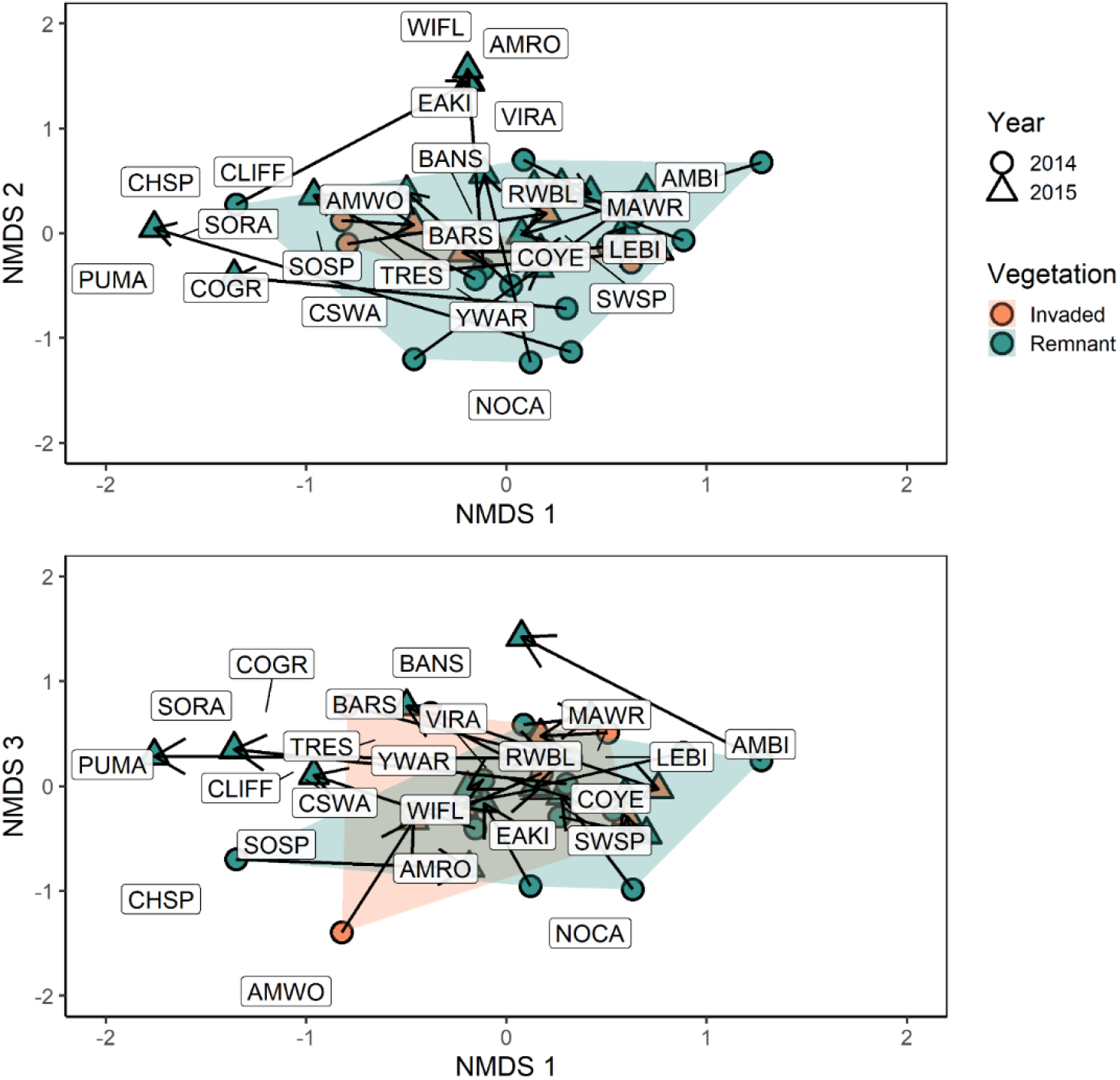
The 3D NMDS ordination solution of Bray-Curtis dissimilarity matrix calculated using bird species abundances (stress = 15.56). The joint plots depict site ordination scores with arrows indicating movement from 2014 to 2015, overlaid with species scores (see Appendix J for codes). Points are surrounded by convex hulls for each vegetation type.

### Beta diversity

Total beta diversity of the bird community was higher in remnant marsh than in *P. australis*-invaded marsh, and beta diversity was mainly driven by bird species turnover in both vegetation communities (Fig. 4; Appendix G). Total beta diversity in *P. australis*-invaded sites was slightly above the expectations of our null model in 2014 and decreased to below expectations in 2015 (Fig. 4). In contrast, beta diversity was stable between years in remnant marsh. Separating remnant vegetation communities into meadow marsh and emergent marsh illustrates how beta diversity was consistently high in emergent marsh both years, while beta diversity went from below the 95% CI of the randomly generated value in 2014 to above the 95% CI in 2015 (Appendix H).

**Figure 4.**
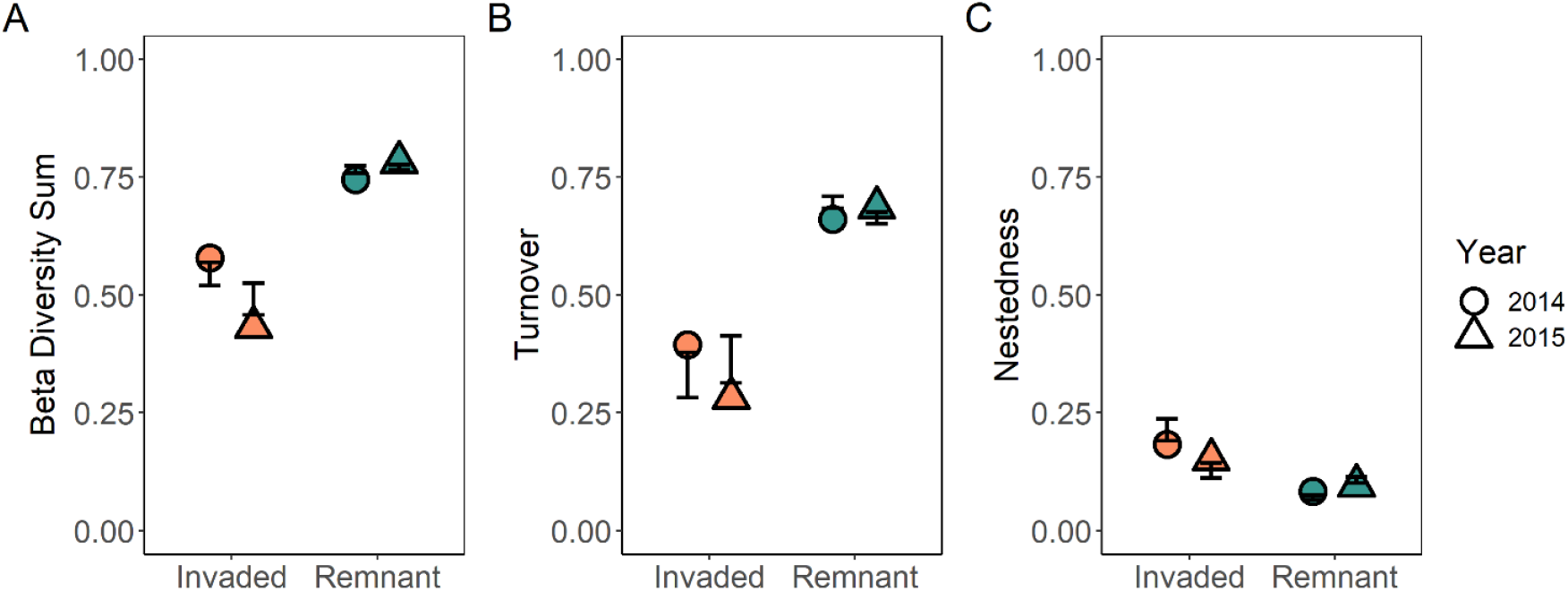
Beta diversity (A), and its turnover (B) and nestedness (C) components calculated using pair-wise Sørensen dissimilarity for remnant marsh vegetation and *P. australis*-invaded marsh. Error bars represent the 95% CI of the null models for vegetation type.

The LCBD results also indicate remnant marsh contributed more to total diversity than *P. australis-*invaded marsh in both years (Fig. 5A & B). In 2014, six remnant emergent marsh sites and one remnant meadow marsh site contributed more than average to diversity (Fig. 5A). In 2015, three meadow marsh and four emergent marsh sites contributed more than average (Fig. 5B). Our comparison of LCBD between years found that half of the remnant meadow marsh sites and two emergent marsh sites became more differentiated in 2015, while the rest of the sites had no change or a slight homogenization (Appendix I). The remnant patches of marsh vegetation had a more unique bird community than the *P. australis-*invaded sites. *Phragmites australis-*invaded sites lost species at four out of six sites, while remnant vegetation sites all gained species in 2015 (Fig. 5C & D; Appendix I).

**Figure 5.**
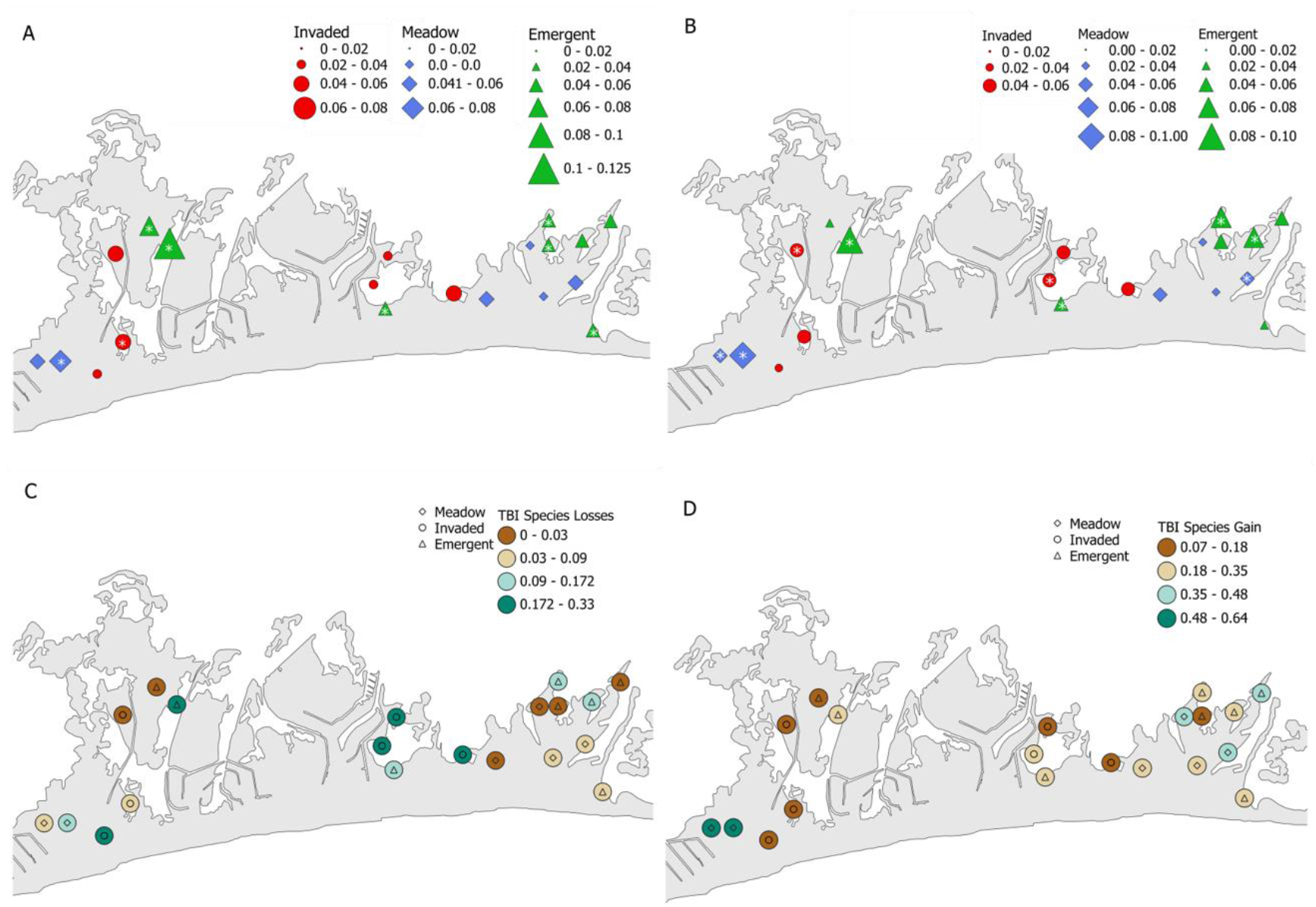
The relative contribution of each site to total beta diversity in 2014 (A) and 2015 (B), where an asterisk (*) indicates sites that contribute more than average. Changes in community composition between years (temporal beta diversity (TBI)) decomposed into species losses (C) and species gains (D) for each site. Figure made with QGIS (QGIS.org (2020) QGIS Geographic Information System. Open Source Geospatial Foundation Project.) and basemap made with Boundary Files, 2016 Census, Statistics Canada Catalogue no. 92-160-X.

## Discussion

Our study provides strong evidence for spatial and temporal homogenization of the wetland bird community in *P. australis*-invaded marsh. *Phragmites australis* replaces native marsh vegetation, resulting in the biotic homogenization of the wetland bird community and the lower contribution of *P. australis*-invaded sites to total dissimilarity. When considering year-to-year changes in the bird communities, the diversity in *P. australis-*invaded sites decreased in response to the deeper water levels in 2015. This is in stark contrast to remnant marsh, and is the opposite trend typically observed between wetland bird communities and deeper standing water (e.g., Timmermans et al. 2008, Jobin et al. 2013, Gnass Giese et al. 2018). Typical measures of alpha diversity failed to detect differences in the bird community between remnant and *P. australis*-invaded marsh. This raises concerns, as such alpha diversity metrics are still widely used to assess biological communities, as noted by Larsen et al. (2018) and Kortz and Magurran (2019). While the average number of bird species in *P. australis*-invaded and remnant marsh were similar, remnant marsh contributed considerably more to gamma and beta diversity. In fact, the bird community present in *P. australis* in both years was mostly composed of a subset of the species using remnant marsh vegetation. Most of the species using *P. australis*, particularly in 2015, were of passerines (Passeriformes), or perching birds, and were also abundant in remnant marsh vegetation. Our results suggest that, while some wetland birds may make use of *P. australis*, this relationship is strongly moderated by differences in average water depths.

The distinctively structured vegetation zones in remnant marsh are likely the reason it supports greater bird diversity. For example, though cattail marsh in deeper water provides tall, robust and fairly dense patches of emergent vegetation similar in canopy height to *P. australis*-invaded patches, in shallower water narrow-leaved emergent vegetation like sedges (Carex. spp.) and Canada Bluejoint grass (*Calamagrostis canadensis*) dominate, creating dense tufts of shorter (ca. 1 m) vegetation with greater floristic diversity, light penetration (Robichaud and Rooney 2021) and distinct litter composition (Yuckin and Rooney 2019), When *P. australis* invades, it displaces both cattail marsh and this shallower meadow marsh zone, as well as the at intermediate water depths, The result is a loss of vegetation diversity, structural complexity and habitat heterogeneity with consequences for avian habitat value. The greater habitat heterogeneity of remnant marsh supports greater taxonomic turnover in the bird community, providing opportunities for numerous bird species to partition resource use within distinct habitats. Habitat heterogeneity through time is also affected. In an average water level year, there were similar numbers of bird species using *P. australis*-invaded marsh and remnant marsh, with fewer species using the dry meadow marsh. However, when water levels increased and meadow marsh flooded, the species richness in *P. australis-*invaded marsh dropped considerably, down to eight of the most commonly observed bird species and the use of meadow marsh by marsh birds increased. Processes, such as invasion, that simplify natural vegetation complexity and reduce habitat complexity through space and time are likely to do harm to wetland-associated bird communities (Steen et al. 2006). We demonstrate that in an average water level year *P. australis* habitat may be used by more species, but when flooding occurs native bird communities preferentially use remnant, uninvaded marsh habitat.

The beta diversity of the bird community in remnant marsh was higher than in *P. australis*-invaded marsh, as we hypothesized, but it was consistent in both years, contrary to what we expected. Examining meadow marsh and emergent marsh as distinct communities provides insight into how the bird community uses remnant marsh. In remnant emergent marsh the bird community was unique and contributed substantially to overall diversity in both years, though beta diversity was consistent between 2014 and 2015. Marsh obligate species such as American Bittern, Least Bittern and Virginia Rail were present in emergent marsh in both study years. The emergent cover, dominated by cattail (*Typha* spp.), and open water interspersion create ideal habitat for these marsh obligates. The permanent standing water in these sites attracts species that consume fish, amphibians, and invertebrates via stalking and probing, and an increase of 20 cm between years likely results in relatively few changes to the resources that emergent marsh adapted species rely on.

Unlike emergent marsh, the bird community using remnant meadow marsh was very responsive to year-to-year water depth differences. In 2015, the number of species using meadow marsh more than doubled and included more marsh obligate species, such as Marsh Wren, Sora, and Virginia Rail, and more aerial insectivore species, like Barn Swallow and Tree Swallow.

Meadow marsh, by nature of its location, represents a transition to upland habitat (Keddy and Campbell 2019). Meadow marsh vegetation provides considerable structural heterogeneity as it contains shrubs growing among hummock-forming grasses and sedges, and patches of flowering forbs. In years when lake water levels are high meadow marsh is also filled with open water pools (Riffell et al. 2001). Many bird species partition their use of similar wetland habitat, such as Common Yellowthroat, which prefer shrubs, and Sora, which prefer open water pools and some floating vegetation (Riffell et al. 2001). The presence of standing water is a cue for some wetland birds to select a breeding territory (Greenberg 1988), and many freshwater invertebrate communities are closely tied to hydrology (Batzer 2013) which may provide more prey in wetter years. Therefore, the change from no standing water to an average of 10 cm is substantial, as depths between 10 – 20 cm provides diverse foraging habitat for wetland birds (e.g., Colwell and Taft 2000). In a year with “average” water depths, the bird community supported by remnant meadow marsh may look less diverse, but our results demonstrate its important contribution to diversity in space and time.

Our research focused on taxonomic homogenization, but it is notable that species loss as a result of invasion is not random and likely leads to functional homogenization (Olden and Rooney 2006). Wetland obligate species American Bittern and Least Bittern were not observed in *P. australis* habitat in either year. In the wetter 2015, the bird community was especially impoverished and only eight of the most abundant bird species were observed using *P. australis* marsh. All eight of these species are passerines and the majority of the species nest in shrubs and consume insects either on the wing or by gleaning from vegetation (Robichaud and Rooney 2017). Wetland obligate species, including rails, bitterns, herons, and other waterbirds, were not present in *P. australis* in 2015. Lockwood et al. (2000) determined that certain families of birds may be more vulnerable to extirpation or extinction as a result of biotic homogenization. They note that Rallidae are at a high risk of extinction – a family of birds that encompasses many wetland species, including Sora and Virginial Rail – in part, due to their specific range requirements (Lockwood et al. 2000). This sheds light on the mechanism by which some wetland species may become extinct: of the IUCN red listed species that are threatened by invasive species, about one third are wetland species (IUCN 2020). Our results indicate that species in the order Passeriformes may fair better in the face of invasive wetland plants, but this requires further study.

While our study focused on only a two-year window in time, it provides compelling evidence that invasive species interact with environmental conditions to change biological communities. The importance of water level fluctuations for wetland bird communities is well documented (e.g., Gnass Giese et al. 2018), but testing the interaction between invasive plants and environmental variables on bird communities presents a challenge. The reported effects of *P. australis* on wetland bird communities in the literature are varied and our results provide direct evidence that water levels mediate *P. australis* use by wetland birds. Importantly, in our study system, *P. australis* is the most abundant invasive plant. The response of birds to invasive plants likely depends on which species is most abundant on the landscape, rather than an innate property of the plant species itself (e.g. Chin et al. 2014). As a model invasive species, however, *P. australis* does possess traits common to other invasive wetland plants such as rhizomatous growth, creating tall dense monocultures, and a wide tolerance to environmental variables (Meyerson et al. 2016). A mechanistic understanding regarding the complex relationship between invasive species and changes to diversity is paramount for adequately conserving and managing ecosystems (e.g., Rooney et al. 2007). This is especially relevant in a time when multiple stressors, including climate change, disturbance, and global connectedness, increase the likelihood of introduced species establishment. Preserving the diversity of wetland habitat is essential to protecting these systems.

## Acknowledgements

Funding for the project was provided by an NSERC Discovery (RGPIN-03846) awarded to R.C.R. We would like to thank Birds Canada for their support with field work, and Heather Polowyk and Graham Howell for their assistance with data collection. We would also like to thank the reviewers whose comments greatly improved the final manuscript.

