## Appendix for "Invasive grass causes biotic homogenization in wetland birds"

Appendices

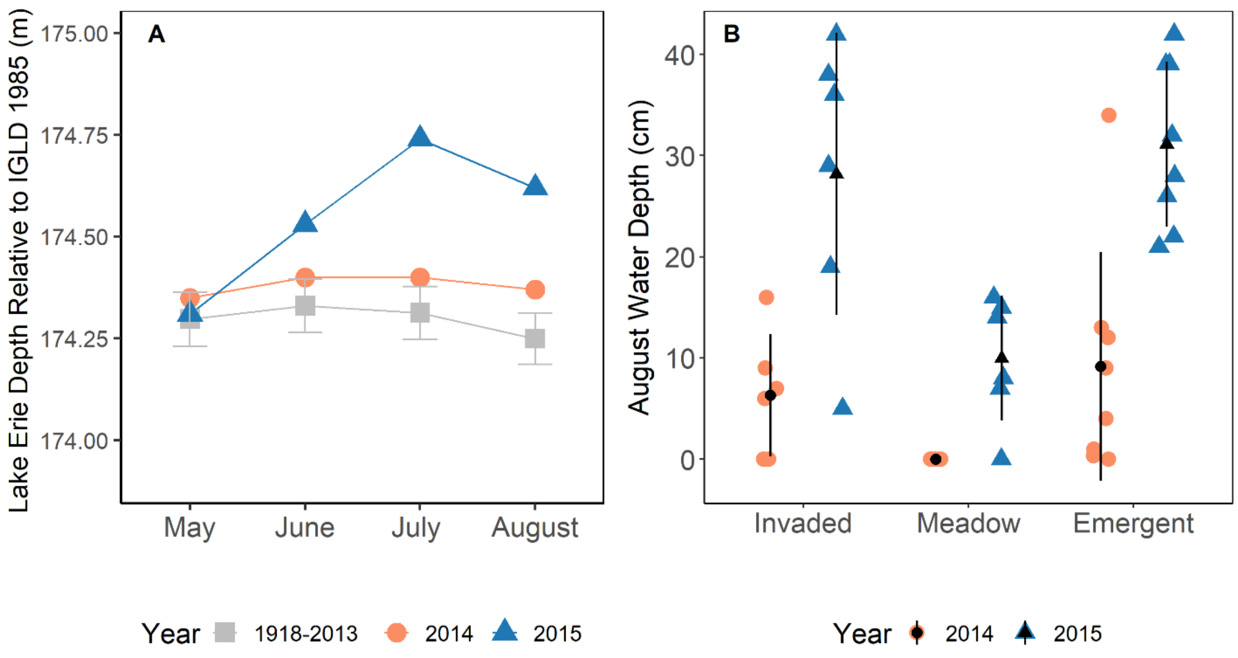


Appendix A. The average monthly Lake Erie water levels during the survey period (A) were slightly above the historical average (grey squares with 95% confidence intervals) in 2014 and considerably higher than average in 2015. Site-level water depths, collected in August, reflected Lake Erie water levels (B). Lake Erie water level averages were gathered from the Canadian Hydrographic Service, interpolated from gauge stations at Port Stanley, Port Colborne, Toledo and Fairport, with depth referenced to the International Great Lakes Datum of 1985 (http://www.tides.gc.ca/C&A/network_means-eng.html, accessed 23 July 2020).

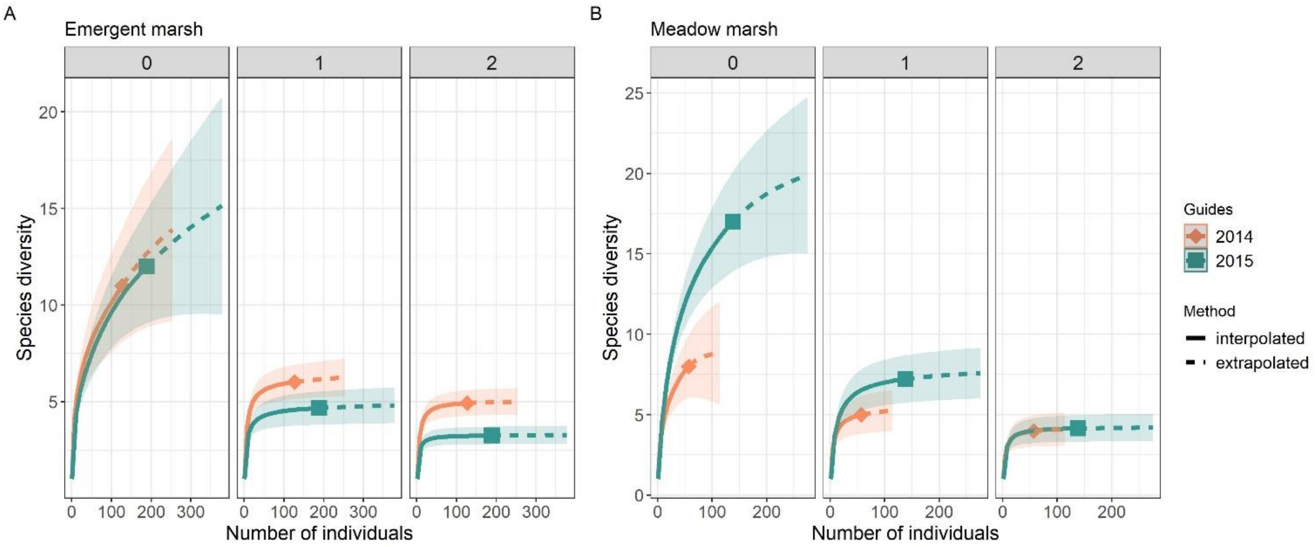


Appendix B. Sample-sized based rarefaction and extrapolation sampling curves for the remnant vegetation, broken into emergent marsh (A) and meadow marsh (B) in 2014 and 2015, by diversity order: 0 = species richness, 1 = Shannon-Weiner diversity, 2 = Simpson's diversity. Points represent the reference samples and shaded areas represent 95% confidence intervals

Appendix C. The total number of each species observed in each vegetation type during the field season.

| Species | Emergent 2014 | Emergent 2015 | Meadow 2014 | Meadow 2015 | *Phragmites* 2014 | *Phragmites* 2015 |
|---|---|---|---|---|---|---|
| American Bittern | 3 | 1 | 0 | 0 | 0 | 0 |
| American Robin | 0 | 0 | 0 | 1 | 0 | 0 |
| American Woodcock | 0 | 0 | 0 | 0 | 1 | 0 |
| Bank Swallow | 0 | 1 | 0 | 1 | 0 | 0 |
| Barn Swallow | 1 | 0 | 0 | 3 | 4 | 2 |
| Chipping Sparrow | 0 | 0 | 1 | 0 | 0 | 0 |
| Cliff Swallow | 0 | 0 | 0 | 1 | 0 | 0 |
| Common Grackle | 0 | 0 | 0 | 1 | 1 | 0 |
| Common Yellowthroat | 24 | 25 | 14 | 19 | 12 | 16 |
| Chestnut-sided Warbler | 1 | 0 | 0 | 1 | 0 | 0 |
| Eastern Kingbird | 0 | 4 | 0 | 5 | 0 | 1 |
| Least Bittern | 1 | 1 | 0 | 0 | 0 | 0 |
| Marsh Wren | 22 | 29 | 0 | 6 | 14 | 15 |
| Northern Cardinal | 0 | 0 | 1 | 0 | 0 | 0 |
| Purple Martin | 0 | 0 | 0 | 2 | 0 | 0 |
| Red-winged Blackbird | 39 | 94 | 22 | 62 | 32 | 34 |
| Sora | 0 | 0 | 0 | 2 | 1 | 0 |
| Song Sparrow | 0 | 0 | 2 | 6 | 2 | 2 |
| Swamp Sparrow | 25 | 25 | 10 | 12 | 19 | 15 |
| Tree Sparrow | 4 | 3 | 2 | 10 | 6 | 0 |
| Virginia Rail | 1 | 3 | 0 | 2 | 1 | 0 |
| Willow Flycatcher | 0 | 1 | 0 | 0 | 0 | 0 |
| Yellow Warbler | 6 | 2 | 5 | 4 | 4 | 4 |

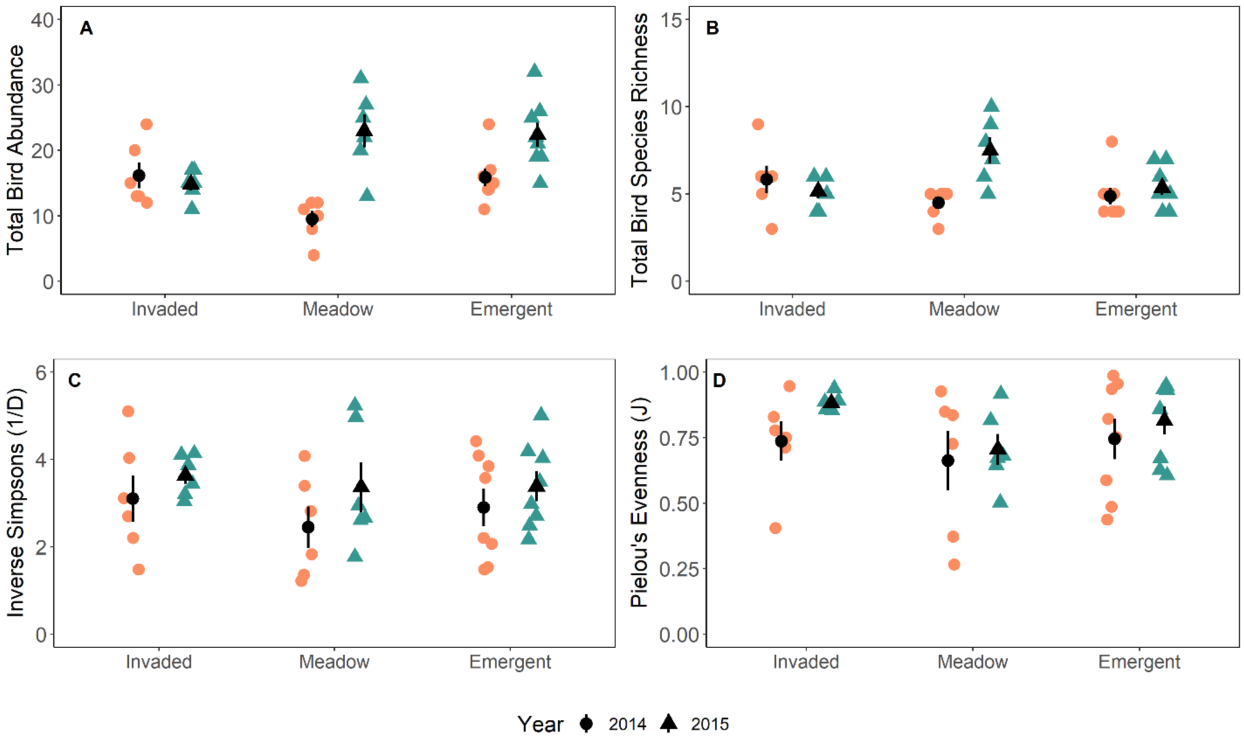


Appendix D. The degree of change in total bird abundance (A), species richness (B), inverse Simpsons (C) and Pielou’s evenness (D) between 2014 and 2015 depends on which vegetation community is considered. Reference meadow and emergent marsh vegetation comprise the “remnant” vegetation community. Lake Erie water levels were higher in 2015, resulting in deeper standing water in all vegetation communities. Error bars represent standard error.

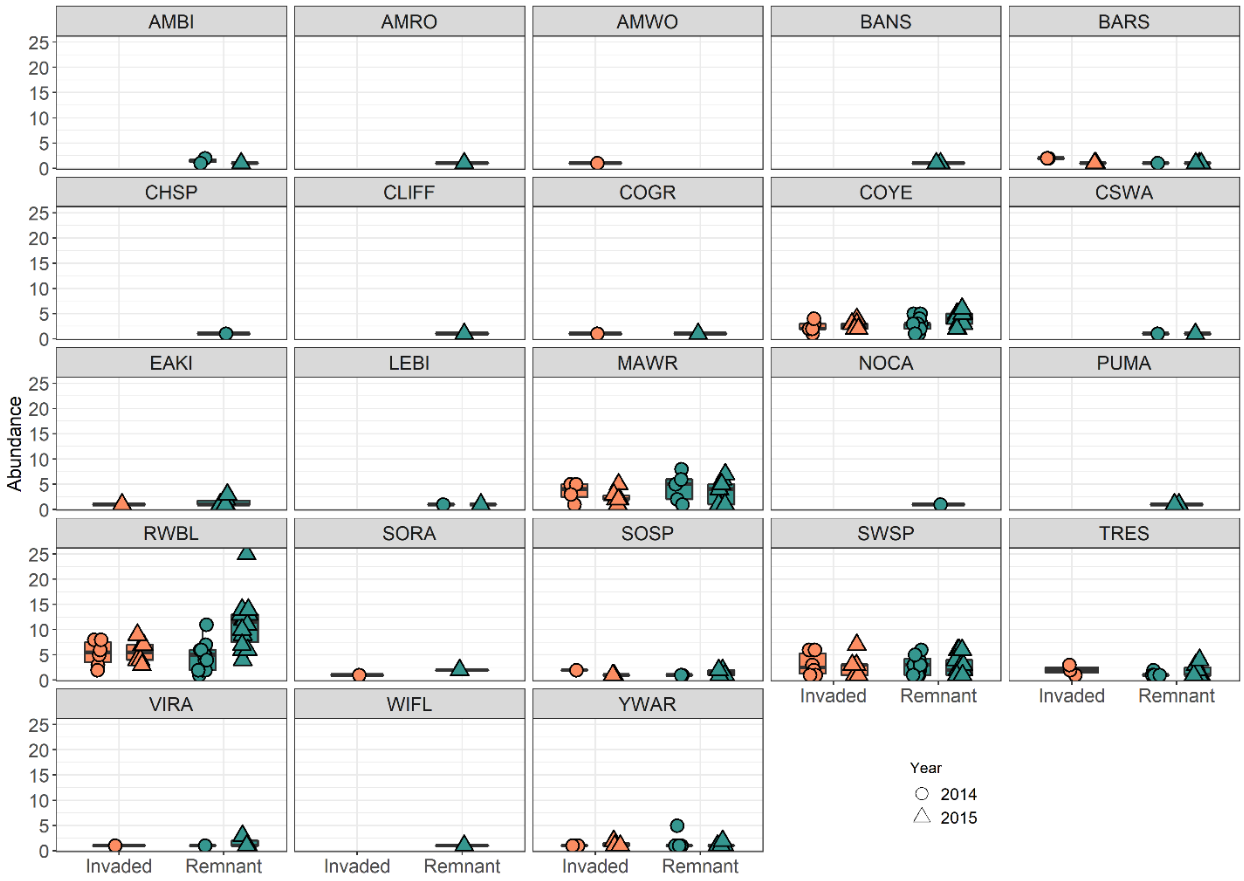


Appendix E. Degree of change for each observed bird species between the average water levels of 2014 and the deeper standing water levels of 2015. Counts of 0 are filtered out to highlight trends between the vegetation types between years. Boxplots represent the 25$^{th}$, 50$^{th}$, and 75$^{th}$ quartile, with whiskers corresponding to 1.5 * IQR and data beyond the whiskers are outliers.

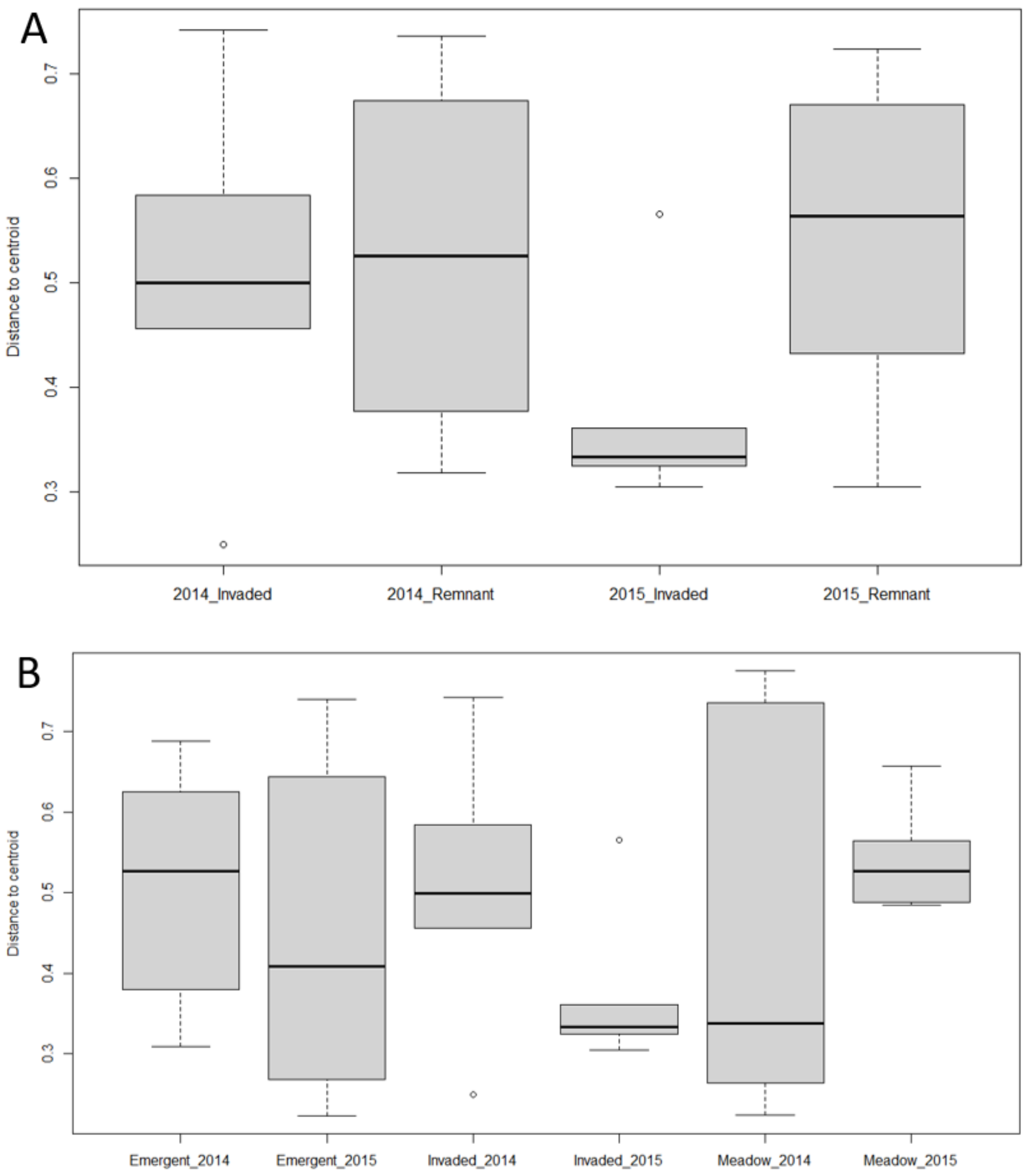


Appendix F. The distance to the group centroid for invaded and remnant marsh in both years (A) and with remnant marsh broken into emergent and meadow marsh (B). Differences were assessed using a pairwise comparisons from the permutational test of homogeneity of multivariate dispersions. Boxplots represent the 25th, 50th, and 75th quartile, with whiskers corresponding to 1.5 * IQR.

Appendix G. Beta diversity, and its nestedness and turnover components, generated using pair-wise Sørensen dissimilarity. Columns include the observed value (“value”) of these metrics for each vegetation type and year combination, and the null model average and 95% CI after 1000 runs with a randomized species matrix.

| | | | Sum | | | Turnover | | | Nestedness | | |
|---|---|---|---|---|---|---|---|---|---|---|---|
| Year | Vegetation | n | Value | Null Average | Null CI | Value | Null Average | Null CI | Value | Null Average | Null CI |
| 2014 | Remnant | 14 | 0.744 | 0.765 | 0.008 | 0.660 | 0.696 | 0.013 | 0.084 | 0.070 | 0.005 |
| 2015 | Remnant | 14 | 0.781 | 0.771 | 0.006 | 0.685 | 0.663 | 0.012 | 0.096 | 0.108 | 0.006 |
| 2014 | Invaded | 6 | 0.578 | 0.544 | 0.025 | 0.395 | 0.330 | 0.048 | 0.183 | 0.214 | 0.023 |
| 2015 | Invaded | 6 | 0.432 | 0.491 | 0.034 | 0.281 | 0.363 | 0.049 | 0.151 | 0.127 | 0.016 |

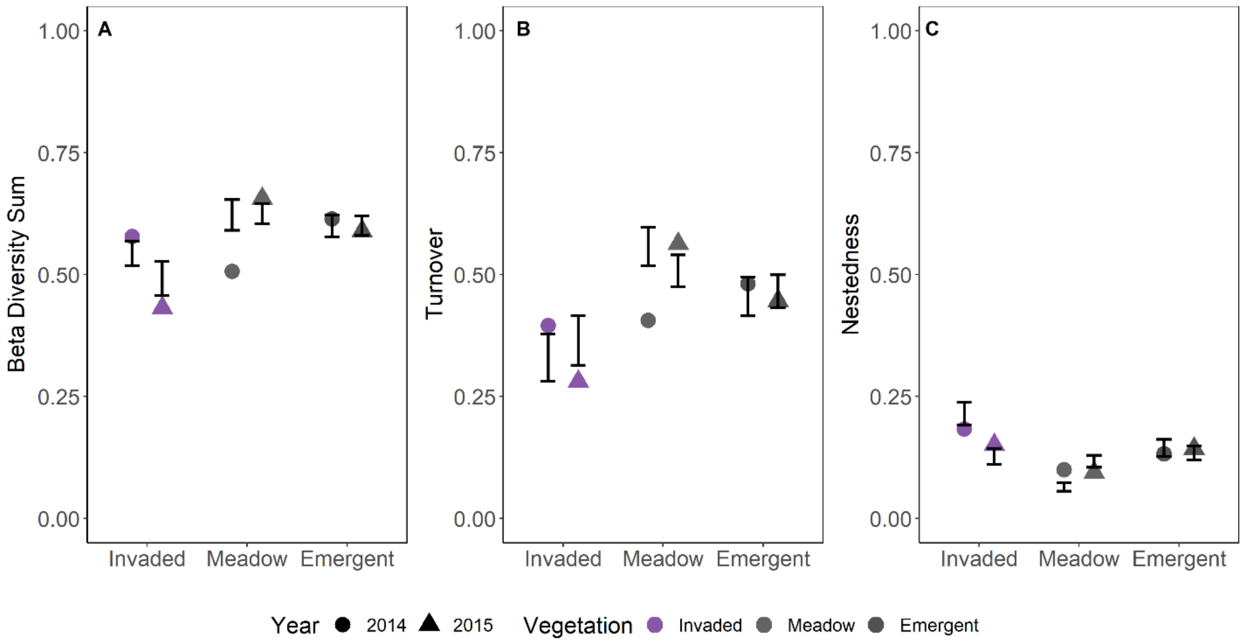


Appendix H. Beta diversity (A), and its turnover (B) and nestedness (C) components, between years for each vegetation type calculated using pair-wise Sørensen dissimilarity. Error bars represent the 95% CI of the null models for vegetation type.

Appendix I. The local contribution to beta diversity (LCBD) of each site in 2014 and 2015, and the difference between the two years ($LCBD_{2015} – LCBD_{2014}$). Temporal beta diversity (TBI) values to assess differences in community composition between both years as total dissimilarity, decomposed into species losses and gains. LCBD and TBI were calculated using Bray-Curtis dissimilarity matrices on raw abundance data.

| | | | LCBD | | | TBI | | | |
|---|---|---|---|---|---|---|---|---|---|
| Site | Vegetation | | 2014 | 2015 | Difference | Loss | Gain | Total | Change |
| CM4R | Emergent | Remnant | 0.121 | 0.095 | -0.027 | 0.241 | 0.276 | 0.517 | + |
| CM5R | Emergent | Remnant | 0.062 | 0.039 | -0.024 | 0.030 | 0.182 | 0.212 | + |
| LP10 | Emergent | Remnant | 0.052 | 0.036 | -0.017 | 0.093 | 0.302 | 0.395 | + |
| LP10R | Emergent | Remnant | 0.058 | 0.057 | -0.001 | 0.162 | 0.351 | 0.514 | + |
| LP15 | Emergent | Remnant | 0.051 | 0.045 | -0.006 | 0.027 | 0.162 | 0.189 | + |
| LP5R | Emergent | Remnant | 0.046 | 0.044 | -0.002 | 0.000 | 0.450 | 0.450 | + |
| LP6R | Emergent | Remnant | 0.042 | 0.069 | 0.028 | 0.098 | 0.317 | 0.415 | + |
| LP7R | Emergent | Remnant | 0.056 | 0.064 | 0.008 | 0.107 | 0.250 | 0.357 | + |
| CM10 | Meadow | Remnant | 0.046 | 0.050 | 0.004 | 0.057 | 0.600 | 0.657 | + |
| CM9 | Meadow | Remnant | 0.075 | 0.090 | 0.015 | 0.118 | 0.647 | 0.765 | + |
| LP12 | Meadow | Remnant | 0.048 | 0.051 | 0.003 | 0.057 | 0.486 | 0.543 | + |
| LP5 | Meadow | Remnant | 0.048 | 0.041 | -0.007 | 0.000 | 0.333 | 0.333 | + |
| LP6 | Meadow | Remnant | 0.028 | 0.026 | -0.002 | 0.094 | 0.344 | 0.438 | + |
| LP8R | Meadow | Remnant | 0.030 | 0.028 | -0.002 | 0.000 | 0.442 | 0.442 | + |
| CM19 | *P. australis* | Invaded | 0.043 | 0.051 | 0.007 | 0.000 | 0.172 | 0.172 | + |
| CM2 | *P. australis* | Invaded | 0.022 | 0.030 | 0.008 | 0.207 | 0.172 | 0.379 | - |
| CM6 | *P. australis* | Invaded | 0.055 | 0.043 | -0.012 | 0.071 | 0.143 | 0.214 | + |
| LP1 | *P. australis* | Invaded | 0.049 | 0.044 | -0.005 | 0.257 | 0.114 | 0.371 | - |
| LP12R | *P. australis* | Invaded | 0.039 | 0.054 | 0.015 | 0.333 | 0.250 | 0.583 | - |
| LP16R | *P. australis* | Invaded | 0.029 | 0.043 | 0.014 | 0.244 | 0.073 | 0.317 | - |

Appendix J. All bird species that were observed throughout the study, including their American Ornithologists' Union common name codes, common name, scientific name, and the Order and Family they belong to.

| Code | Common Name | Scientific Name | Order | Family |
|---|---|---|---|---|
| AMBI | American Bittern | *Botaurus lentiginosus* | Pelecaniformes | Ardeidae |
| AMRO | American Robin | *Turdus migratorius* | Passeriformes | Turdidae |
| AMWO | American Woodcock | *Scolopax minor* | Charadriiformes | Scolopacidae |
| BANS | Bank Swallow | *Riparia riparia* | Passeriformes | Hirundinidae |
| BARS | Barn Swallow | *Hirundo rustica* | Passeriformes | Hirundinidae |
| CHSP | Chipping Sparrow | *Spizella passerina* | Passeriformes | Emberizidae |
| CLIFF | Cliff Swallow | *Petrochelidon pyrrhonota* | Passeriformes | Hirundinidae |
| COGR | Common Grackle | *Quiscalus quiscula* | Passeriformes | Icteridae |
| COYE | Common Yellowthroat | *Geothlypis trichas* | Passeriformes | Parulidae |
| CSWA | Chestnut-sided Warbler | *Setophaga pensylvanica* | Passeriformes | Parulidae |
| EAKI | Eastern Kingbird | *Tyrannus tyrannus* | Passeriformes | Tyrannidae |
| LEBI | Least Bittern | *Ixobrychus exilis* | Pelecaniformes | Ardeidae |
| MAWR | Marsh Wren | *Cistothorus palustris* | Passeriformes | Troglodytidae |
| NOCA | Northern Cardinal | *Cardinalis cardinalis* | Passeriformes | Cardinalidae |
| PUMA | Purple Martin | *Progne subis* | Passeriformes | Hirundinidae |
| RWBL | Red-winged Blackbird | *Agelaius phoeniceus* | Passeriformes | Icteridae |
| SORA | Sora | *Porzana carolina* | Gruiformes | Rallidae |
| SOSP | Song Sparrow | *Melospiza melodia* | Passeriformes | Emberizidae |
| SWSP | Swamp Sparrow | *Melospiza georgiana* | Passeriformes | Emberizidae |
| TRES | Tree Swallow | *Tachycineta bicolor* | Passeriformes | Hirundinidae |
| VIRA | Virginia Rail | *Rallus limicola* | Gruiformes | Rallidae |
| WIFL | Willow Flycatcher | *Empidonax traillii* | Passeriformes | Tyrannidae |
| YWAR | Yellow Warbler | *Setophaga petechia* | Passeriformes | Parulidae |
